# Setaphyte VERY-LONG-CHAIN FATTY ACYL DESATURASES are desaturases that impact glycerolipid and sphingolipid metabolism

**DOI:** 10.1101/2025.08.12.669868

**Authors:** Pauline Duminil, Cornelia Herrfurth, Tegan M. Haslam, Ivo Feussner

## Abstract

Desaturases in plants are diverse. They vary in localization, source of reducing power, and substrate preference, accepting glycerolipids, long-chain bases, acyl-CoAs, and acyl-ACPs, in varying states of (un)saturation and chain length. Their products are incorporated into membrane glycerolipids, sphingolipids, or storage lipids. We previously characterized a desaturase from *Physcomitrium* patens that predominantly affects the monounsaturation of very-long-chain fatty acyl (VLCFA) moieties of sphingolipids, naming this desaturase SPHINGOLIPID FATTY ACYL DESATURASE (SFD). Among embryophytes, candidate SFDs were only identified in setaphytes, including one paralog in *P. patens* and an ortholog in *Marchantia polymorpha*. Here, we characterize the *P. patens* paralog, and clarify via mutant analysis that SFDs affect not only sphingolipid metabolism, but also glycerolipid metabolism. We express both paralogs, as well as the candidate gene from *M. polymorpha*, in *Saccharomyces cerevisiae*, and show they desaturate VLCFAs incorporated into sphingolipids, triacylglycerols, and acyl-CoAs. The simplest explanation is that “SFDs” likely accept an acyl-CoA, rather than a sphingolipid substrate, as initially proposed. We suggest renaming these desaturases VERY-LONG-CHAIN FATTY ACYL DESATURASES (VFADs). The physiological functions of VFADs and analogous enzymes from other plant systems are discussed, as are the challenges with classifying desaturase activities.

**Highlight:** VFADs from *P. patens* and *M. polymorpha* produce monounsaturated very-long-chain fatty acids that are incorporated into sphingolipids and glycerolipids, likely via desaturation of an acyl-CoA substrate

## Introduction

Fatty acyl desaturation is executed by a broad diversity of enzymes with wide-ranging metabolic roles, both within and among different plant lineages. Enzymes with diverse structures and phylogenetic origins contribute to acyl desaturation. Two major desaturase groups are soluble acyl-acyl carrier protein (ACP) desaturases, and transmembrane histidine box desaturases (Shanklin and Cahoon, 1998; Kazaz *et al*., 2022). There is further diversity among transmembrane desaturases based on substrate identity, that is, acyl linkage to Coenzyme A (CoA), a glycoglycerolipid, a phosphoglycerolipid, or a sphingolipid, and additionally the acyl chain length and existing saturation status. The relative position of the introduced desaturation from either the methyl or carboxyl end of the lipid, or to existing unsaturated bonds, also contributes to the diversity of acyl products (Napier *et al*., 1999; Sperling *et al*., 2003)

In plants, soluble, plastidial acyl-ACP desaturases are responsible for a first desaturation of *de novo* synthesized long-chain fatty acids (LCFAs) to produce monounsaturated LC-acyl-ACPs. These are the best-characterized plant desaturases, as their solubility facilitates biochemical work that is difficult with the remaining integral membrane desaturases. While there is diversity in the position of this initial desaturation, in *A. thaliana* and in many other plants the first desaturation produces 18:1^Δ9^ (Shanklin and Somerville, 1991). Subsequently, a series of acyl-lipid desaturases (FATTY ACID DESATURASEs, FADs, identified in mutant screens of *A. thaliana*, reviewed in (Ohlrogge and Browse, 2011), further desaturate LCFAs esterified to different glycerolipid substrates, depending on the enzyme localization. Two of these, FAD4 and FAD5, accept saturated substrates to produce monounsaturated FAs within the plastid. FAD4 produces 16:1^Δ3E^ on phosphatidylglycerol (PG) that is unique in its position and stereochemistry, and the 16:1^Δ3E^ product remains unique to PG (Browse *et al*., 1985). FAD5 produces 16:1^Δ7^ on monogalactosyldiacylglycerol (MGDG), which can be further desaturated and also transferred to other lipids (Kunst *et al*., 1989). The remaining FAD acyl-lipid desaturases are methyl-end desaturases that accept unsaturated substrate (initially either the 18:1^Δ9^ product of the acyl-ACP desaturase, or 16:1^Δ7^ product of FAD5) and introduce additional double bonds between this and the methyl end of the acyl group. In plastids FAD6 accepts 16:1- and 18:1-esterified plastidial glycerolipids to produce 16:2 and 18:2 ω-6 fatty acids (Browse *et al*., 1989), and FAD7 and FAD8 redundantly desaturate 16:2 and 18:2 to 16:3 and 18:3 ω-3 fatty acids, respectively, as components of the same plastidial glycerolipids (Browse *et al*., 1986; Gibson *et al*., 1994; McConn *et al*., 1994). In the endoplasmic reticulum (ER), FAD2 desaturates 18:1 (18:1^Δ9^ from the acyl-ACP desaturase) to 18:2^Δ9,12^, and FAD3 desaturates 18:2^Δ9,12^ to 18:3^Δ9,12,15^ (Arondel *et al*., 1992; Miquel and Browse, 1992; Browse *et al*., 1993).

One facet of desaturase enzyme diversification among plants specifically concerns the synthesis of monounsaturated very-long-chain fatty acids (VLCFAs), occurring at the ER, and distinctly from plastidial acyl-ACP desaturase, FAD4, and FAD5 activity. Monounsaturated VLCFAs are primarily used in the synthesis of glycosyl inositol phosphorylceramides (GIPCs), a class of plasma membrane (PM)-enriched sphingolipids, as well as some lower-abundant membrane glycerolipids. In the model tracheophyte *A. thaliana*, this desaturation was shown to be executed by ACYL-COENZYME A DESATURASE (ADS) LIKE 2 (AtADS2), a member of a desaturase family with homology to yeast and mammalian acyl-CoA desaturases and acyl-lipid desaturases of cyanobacteria (Smith *et al*., 2013). Plant ADSs generally accept acyl-CoA substrates; however, this is not an absolute rule, as there is evidence that AtADS2 could alternatively accept a galactolipid substrate (Heilmann *et al*., 2004; Chen and Thelen, 2013). Further diversity within the ADS gene family arises from the fact that, based on their activity with 24:0 and 26:0 substrates in yeast, different members of the family can be considered to establish their position for monounsaturation relative to either the carboxyl (AtADS1.2 and AtADS1.4) or methyl (AtADS1, AtADS1.3, AtADS2, AtADS4, and AtADS4.2) end of the acyl group (Smith *et al*., 2013). In spite of these complexities, conserved sequence features such as the absence of an N-terminally fused cytochrome b5 domain and conserved histidine boxes (discussed below) support the clustering of this group (Shanklin and Cahoon, 1998; Sperling *et al*., 2003).

We recently discovered that in the model bryophyte *Physcomitrium patens* (formerly *Physcomitrella patens*), the monounsaturation of VLCFA moieties of sphingolipids is catalyzed by SPHINGOLIPID FATTY ACYL DESATURASE (PpSFD, gene identifiers V1.6: Pp1s286_53V6.1, V3.3: Pp3c21_180V3.1, V6: Pp6c21_120V6.1), a cytochrome-b5 fusion desaturase with homology to long-chain base desaturases and the front-end desaturases characteristic of algal and bryophyte long-chain polyunsaturated lipid synthesis (Resemann *et al*., 2021). SFD is unlike ADSs in domain structure, phylogeny, and expected substrate preference. Classification as a methyl-end (as AtADS2) or a carboxyl-end (as PpSFD) desaturase is in itself a marked difference. In connection to this, carboxyl-end desaturases characteristically include an N-terminally-fused cytochrome b5 domain that confers reducing power to the enzyme, whereas methyl-end desaturases require an external electron donor: cytochrome b5 is used in the case of microsomal desaturases, and ferredoxin in the case of plastidial desaturases (Sperling and Heinz, 2001). The catalytic histidine box structure of the two enzyme classes is also considered a distinguishing feature; the position of the first histidine residue within the third box of carboxyl-end desaturases is replaced by a glutamine, that is, Q(X)_2-3_HH replaces H(X)_2-3_HH.

In spite of these differences, the physiological functions of AtADS2 and PpSFD enzymes are convergent and similarly contribute to cold stress resilience, to the extent that heterologous expression of *PpSFD* in *A. thaliana* can complement the cold-sensitivity of *Atads2* knock-out mutants (Resemann *et al*., 2021). While increased desaturation of membrane lipids is a ubiquitous and extensively-reported adaptation to cold stress (Chen and Thelen, 2013), this specific modification impacting the same product sphingolipids, catalyzed by such seemingly different enzymes, is striking and unique.

We previously reported that knockout *Ppsfd* mutants had depleted, but not completely abolished levels of monounsaturated VLCFAs in both sphingolipids and phosphoglycerolipids, indicating that another gene in the *P. patens* genome must contribute to this metabolic role. A clear candidate that could produce monounsaturated VLCFAs in *P. patens* is SFD-LIKE (PpSFD-LIKE, gene identifiers V1.6: Pp1s3_40V6.1, V3.3: Pp3c18_17160V3.1, V6: Pp6c18_7810V6.1), a gene product with 79 % amino acid identity to SFD, and a full complement of the histidine boxes required for catalysis (Supplemental Figure 1). According to public microarray and RNAseq data, *PpSFD-LIKE* expression levels are consistently lower than *PpSFD*; however, *PpSFD-LIKE* does have elevated expression levels in skotonema, a specialized type of filamentous protonemal tissue produced in the dark (Ortiz-Ramírez *et al*., 2016; Fernandez-Pozo *et al*., 2020). We speculated that *PpSFD-LIKE* may support the metabolic role of *PpSFD*, and have a physiological function that is important under particular environmental stresses. To explore this, we generated *Ppsfd-like* single and *Ppsfd sfd-like* double mutants by CRISPR/Cas9 mutagenesis, and performed lipidomic profiling of the mutants grown under standard and dark conditions. We were also curious about the metabolic functions of the *SFD-LIKE* gene family more broadly, and skeptical of the initial assessment of SFD-LIKEs as sphingolipid desaturases that implied they accept ceramide substrates. We therefore measured glycerolipids of the mutants, and expressed *PpSFD* and other *SFD-LIKEs* in yeast. We then profiled the sphingolipids, acyl-CoAs, and VLCFA-accumulating glycerolipids in the yeast cell lines. Altogether our results suggest that SFD-LIKE activity is conserved in setaphytes, and that an acyl-CoA substrate of SFDs presents the most harmonious model for SFD functions in lipid metabolism. In light of these results, we suggest renaming SFDs as VERY-LONG-CHAIN FATTY ACYL DESATURASEs (VFADs), with *PpSFD* becoming *PpVFAD1*, and *PpSFD-LIKE* becoming *PpVFAD2*. This nomenclature maintains the distinction from ADSs and from FADs, and highlights the clear and consistent feature of these enzymes accepting very-long-chain substrates. Characterization of *Ppvfad1 vfad2* double mutants lacking monounsaturated VLCFAs demonstrates that these are dispensable for normal growth, though they contribute to fitness under cold stress conditions. We discuss the phylogeny and specificity of desaturase activity, and possible questions to be explored next.

## Materials and methods

### *P. patens* culture conditions

The wild-type strain Gransden 2004 and was obtained from the international Moss Stock Center (IMSC; https://www.moss-stock-center.org/en/strain#40001), hereafter referred to simply as the wild type, or WT. The *sfd10* (hereafter *vfad1*) mutant was previously generated and described by Resemann *et al*., 2021. Unless otherwise indicated, plants were cultivated under long day conditions 16 h light/8 h dark with a light intensity of 105-120 μmol m^-2^ s^-1^ at 25 °C/18 °C, in 9 cm petri dishes. Protonema was propagated every 7-10 days by blending tissue in sterile tap water with an ULTRA-TURRAX® (IKA) for 20 s. The resulting cell suspension was transferred onto BCD-AT (Maronova and Kalyna, 2016) medium overlaid with sterile cellophane discs. Protonema was used as starting material to produce skotonema (Saavedra *et al*., 2015); spot inocula were transferred to square petri dishes containing BCD-AT medium with 2% sucrose and grown for 7 days. The plates were then positioned vertically in the dark for 4 weeks before harvesting for lipid analysis. To produce gametophores, protonema spot inocula were placed on petri dishes containing BCD medium for 4-5 weeks.

For cold treatment, petri dishes were sealed with parafilm and transferred to a chamber at 6°C with 55–70 μmol m^−2^ s^−1^ continuous light in order to reproduce the same stress as in Resemann *et al*., 2021.

### Gene editing *via* CRISPR-Cas9

To obtain CRISPR-Cas9 mutants of *sfd-like*, hereafter *vfad2*, CRISPR single guides were designed using CRISPOR (http://crispor.tefor.net/) and synthetized by Merck as oligonucleotides. Mixtures of the forward and reverse oligos encoding each guide were phosphorylated with T4 phosphonucleotide kinase (PNK) (NEB), the enzyme was subsequently heat inactivated, and the oligos were allowed to anneal into duplexes by slowly cooling the reaction mixture. The pUCRISPR vector (Haslam *et al*., 2024) was linearized with BbsI (Fermentas) and dephosphorylated using FastAP thermosensitive alkaline phosphatase (Thermo Fisher Scientific). Plasmids expressing the Cas9 nuclease (pAct1-Cas9) and G418 selection marker (pBNRF) were provided by Prof. Fabien Nogué (INRA Versailles) (Lopez-Obando *et al*., 2016; Collonnier *et al*., 2017). Both wild type and *vfad1* were transformed in order to generate single *vfad2* and double *vfad1 vfad2* mutants, respectively.

### *P. patens* protoplast transformation

The protoplast preparation and PEG-mediated transformation of the moss *P. patens* was adapted from (Liu and Vidali, 2011). One-week-old protonema tissue was incubated for approximately 3 h in a solution of 1 % (w/v) Driselase and 8 % (w/v) mannitol. The cells were filtered through a 100 µm sterile mesh and washed twice by centrifugation (Centrifuge 5810R, Eppendorf) at 130 g for 5 min using 15 mL 8 % (w/v) mannitol. After each centrifugation, the supernatant was removed and after the last washing cycle, the cells were resuspended in 2-3 mL mannitol magnesium solution (0.4 M mannitol, 15 mM MgCl_2_, 4mM MES, pH 5.7) according the cell concentration, and incubated for 30 min at room temperature. 600 µL of the cell suspension was used for each individual transformation, by mixing 10 µg pAct1-Cas9, and equal amounts of pBNRF and the single guide constructs in pUCRISPR. 700 µL PEG-calcium solution (4 g PEG4000, 3 mL H_2_O, 2.5 mL 0.8 M mannitol, 1 mL 1 M CaCl_2_) was also added to the cell and plasmid suspension before another 30 min incubation at room temperature. PRM-T medium (modified BCD-AT medium with 10 mM CaCl_2_, 6 % mannitol and 0.4 % plant agar) was melted and kept molten at 42°C. After the incubation period, the protoplast suspension was diluted with 3 mL W5 buffer (154 mM NaCl, 125 mM CaCl_2_, 5 mM KCl, 2 mM MES, pH5.7), the protoplasts were centrifuged at 130 g, and the supernatant was removed. The protoplasts were finally resuspended in 5 mL PRM-T and rapidly distributed onto PRM-B medium (modified BCD-AT medium with 10 mM CaCl_2_ and 6 % (w/v) mannitol) overlaid with cellophane. Five days after transformation, resistant plants were selected on 40 mg/L G418, and subsequently individually transferred to BCD medium plates for further growth and genotyping. Genomic DNA was extracted using the Extract-N-Amp™ Plant Tissue Kits (Sigma) according to the manufacturer’s instructions. The region of interest (approximately 600 bp) surrounding the targeted sequence was amplified by PCR and cleaned up for DNA sequencing (Eurofins). All primers used for amplification and sequencing are listed in Supporting Table S1. Two different guides were cloned but one was more efficient, therefore the *vfad2* mutants used in this study have different lesions at the same target site.

### RNA extraction, cDNA synthesis, and qPCR

RNA was extracted from 10 mg of lyophilized material using the Spectrum™ Plant Total RNA Kit (Sigma) according to the manufacturer’s instructions, and treated with DNAse I prior to cDNA synthesis. Each sample represents three separately grown, harvested, and processed biological replicates. RNA concentrations were measured with a Nanodrop© spectrophotometer, and 1 µg of the treated RNA was used to synthesize cDNA using the RevertAid Minus Reverse Transcriptase (Thermo Fisher Scientific) according to the manufacturer’s instructions.

cDNA from protonema, gametophores, and skotonema were used to measure *VFAD2* expression with the Takyon™ No Rox SYBR® MasterMix dTTP Blue (Eurogentec) according to the manufacturer’s instructions and an iQTM5 PCR as detection system. A previously-tested E2 ubiquitin-conjugating enzyme (Pp6c14_11780V6.1) was used as housekeeping gene (Le Bail *et al*., 2013). All the primer combinations used are described in Supporting Table S1. Primer efficiency for two pairs specific to *PpVFAD2* were tested using a range of cDNA concentrations from 0 to 100 ng. The most efficient primer pair (94.11 %) was used in the presented experiments. Melt-curve analysis were also performed as quality control upon increased temperature from 60°C to 95°C.

### Heterologous expression in the *S. cerevisiae* mutant *ole1*

Sequences of VFAD (“SFD”) proteins were identified and their phylogeny was described in Resemann *et al*., 2021. The coding sequences were retrieved from Phytozome (https://phytozome-next.jgi.doe.gov/), and amplified from cDNA synthesized from RNA extracted from *P. patens* gametophores, *Marchantia polymorpha* thalli (kindly provided by Dr. Alisa Keyl), and *Thalassiosira pseudonana* (Schwarz *et al*., 2022) and inserted in pYES2 (Thermo Fisher Scientific) with uracil selection. Cloning primers and gene identifiers are listed in Supplemental Table S1. The plasmids were verified by colony PCR and Sanger sequencing. A construct for expression of *Chlamydomonas reinhardtii VFAD-LIKE* (Cre10.g453600) was obtained by gene synthesis (Biocat) with codon optimization in a pUC18 vector, and subcloned into pYES2.

Competent *oleic acid requiring* (*ole1*) mutant cells of *S. cerevisiae* (Stukey *et al*., 1990) were then transformed with 5 µg of each of the following plasmids: empty vector pYES2, pYES2-PpVFAD1 (Resemann *et al*., 2021), pYES2-PpVFAD2, pYES2-MpVFAD-LIKE, pYES2-CrVFAD-LIKE, pYES2-TpVFAD-LIKE1, pYES2-TpVFAD-LIKE2. Transformed colonies were selected on synthetic complete medium lacking uracil (SC-U), and confirmed by colony PCR. The cultivation was adapted slightly from Resemann *et al*., 2021. Three to four individual colonies were grown overnight in 5 mL pre-cultures in SC-U, 2 % (w/v) raffinose, and 1 mM linoleic acid to rescue the *ole1* mutant lacking unsaturated fatty acids. Equivalent optical densities of cells of each genotype were used to inoculate 20 mL cultures containing SC-U with 2 % (w/v) galactose and 1 mM linoleic acid. These were grown for 5 days at 28°C, and then harvested by centrifugation at 1000 g for 20 minutes, washed once with water, and stored at -80°C until lyophilization.

### Lipid extraction*s* from *P. patens* and *S. cerevisiae*

Lipids were extracted from lyophilized yeast or plant material according to Herrfurth *et al*., 2021. Lyophilized plant material was crushed with metal beads and the powder resuspended in 6 mL extraction buffer (propan-2-ol/hexane/water (60:26:14, v/v/v)). Lyophilized yeast material was directly resuspended in 6 mL extraction buffer together with glass beads. For both tissue types, the suspended material was incubated at 60 °C for 30 min, and every 10 min, the samples were sonicated for 1 min and briefly vortexed. The cell debris was spun down by centrifugation (Centrifuge 5810R, Eppendorf) for 20 min at 800 g, the supernatant was transferred to a fresh tube, and the solvents were dried under a nitrogen stream at 40 °C. The lipids were finally resuspended in 800 μL of tetrahydrofuran/methanol/water (4:4:1, v/v/v), covered with argon gas, and kept at -20 °C.

### Lipidomic analysis

The multiple reaction monitoring (MRM)-based ultraperformance liquid chromatography nano-electrospray ionization tandem mass spectrometry (UPLC–nanoESI–MS/MS) method described in Tarazona *et al*., 2015 was used to identify and quantify different lipid species. 2 µL of each sample was injected at 18 °C, and lipids were separated *via* reversed-phase UPLC using an ACQUITY UPLC I-class system (Waters) with an ACQUITY UPLC HSS T3 column (100 mm × 1 mm, 1 μm; Waters), at a flow rate of 0.1 mL/min. Solvents were (solvent A) methanol:20 mM ammonium acetate (3:7, v/v) with 0.1 % (v/v) acetic acid, and (solvent B) tetrahydrofuran:methanol:20 mM ammonium acetate (6:3:1, v/v/v) with 0.1 % (v/v) acetic acid. The same gradient (gradient 2b) was used for separating all phosphoglycerolipids and all sphingolipids. For neutral lipids, the flow rate was increased to 0.13 mL/min and the gradient was also adjusted (gradient 1) (Herrfurth *et al*., 2021).

An AB Sciex 6500 QTRAP tandem mass spectrometer was used for analysis. For ceramides and hexosylceramides, precursor ions were [M+H]^+^ and product ions were dehydrated LCBs. For GIPCs, inositol phosphorylceramides (IPCs), mannosyl inositol phosphorylceramides (MIPCs), and mannosyl diinositol phosphorylceramides (M(IP)_2_ Cs), precursor ions were [M+H]^+^ and product ions were ceramides. For the phosphoglycerolipids phosphatidylcholine (PC), phosphatidylethanolamine (PE), and phosphatidylinositol (PI), precursor ions were [M]^-^ and product ions were fatty acids. For triacylglycerols (TAGs), the precursor ions were [M+H]^+^ and product ions were fatty acids.

### Fatty acid methyl ester (FAMEs) derivatization and analysis

Fatty acids from 40 µL of the lipid extracts were methyl esterified using sulfuric acid solution (Garcés and Mancha, 1993). 5 μg tripentadecanoin standard was added to each sample, the solvents evaporated, and residue suspended in 1 mL sulfuric acid solution. The methanolysis proceeded in an 80 °C water bath for 1 h. The reaction was stopped by addition of 200 µL saturated NaCl, and the FAMEs were extracted in 1 mL hexane. The hexane (upper phase) was siphoned off, dried, and the FAMEs residues finally resuspended in 10 µL acetonitrile to be injected into an Agilent 6890 gas chromatography-flame ionization (GC-FID) system with a DB-23 column (30 m x 0.25 mm, 0.25 µm coating thickness, Agilent). Helium was used as carrier gas at a flow rate of 1 ml/min. The temperature gradient was 150 °C for 1 min, 150 – 200 °C at 4 °C /min, 200–250 °C at 5 °C /min and 250 °C for 6 min.

### Acyl-CoA extraction from *S. cerevisiae*

Acyl-CoA extraction protocol was adapted from (Larson and Graham, 2001; Larson *et al*., 2002; Domergue *et al*., 2005). Briefly, after lyophilization, 60 mg of *S. cerevisiae ole1* cells were crushed using stainless steel beads for 30 seconds. Then, 200 µL of freshly prepared ice-cold extraction buffer (1 mL isopropanol, 1 mL 50 mM KH_2_ PO_4_ pH7.5, 40 µL 50 mg mL^−1^ fatty acid-free BSA (stored at -20 °C), and 25 µL glacial acetic acid) was added to the material, followed by the addition of 20 µL 2 pmol µL^-1^ 17:0-CoA as internal standard. The mixture was carefully ground using an Eppendorf pestle. Lipids and pigments were removed through four washing steps using 300 µL of warm petroleum ether saturated using isopropanol/water (1:1, v/v) kept at 50 °C. The mixture was briefly vortexed, then centrifuged at 400 g for 1 min, and the upper phase carefully removed and discarded. The aqueous phase was then transferred to a clean Eppendorf tube and 5 µL of saturated (NH_4_)_2_SO_4_ and 600 µL MeOH/CHCl_3_ (2:1, v/v) were successively added before vortexing and incubating for 20 min at room temperature. After a short centrifugation step (16,000 g for 2 min) the upper phase was transferred to a sample vial for a drying step at 50 °C using a nitrogen evaporator. The dried extract was resuspended in 40 µL of H_2_0/acetonitrile (9:1, v/v) containing 15 mM ammonium hydroxide for analysis.

### Determination of Acyl-CoAs by HPLC-ESI-MS/MS

The analysis of acyl-CoAs was performed accordingly to Haslam and Larson, 2021, with some modifications. Analysis of constituents was achieved using an Agilent 1100 high performance liquid chromatography (HPLC) system equipped with an Agilent ZORBAX RR Eclipse XDB-C18 column (100 mm x 2.1 mm, 3.5 µm) and a quaternary pump and coupled to an AB Sciex 4000 QTRAP tandem mass spectrometer. Aliquots of 35 µl were injected and a ternary gradient of eluent A (acetonitrile, 15 mM ammonium hydroxide), eluent B (acetonitrile/water, 9:1, v/v, 15 mM ammonium hydroxide) and eluent C (acetonitrile/water, 7:3, v/v, containing 0.1 %, v/v formic acid) was applied as follows: 0 min 100 % B, 0-5 min linear gradient to 25 % A and 75 % B, 5-11 min linear gradient to 100 % A, 11-13 min 100 % A, 13-15 min linear gradient to 100 % C, 15-18 min 100 % C, 18-20 min linear gradient to 100 % B and 20-30 min 100 % B. The flow rate was 0.2 ml/min and the separation temperature was constantly at 30 °C.

Tandem MS analysis was performed with a turbo V electrospray ionization source. Ion spray voltage was set to 5.5 kV, source temperature to 300 °C, ion source gas 1 and 2 were set at 60 (arbitrary units) and curtain gas was set at 30 (arbitrary units). Acyl-CoAs were positively ionized and determined in multiple reaction monitoring mode. Mass transitions were as follows: 1118.6/611.6 [declustering potential (DP) 180 V, entrance potential (EP) 10 V, collision energy (CE) 60 V] for 24:0-CoA, 1116.6/609.6 (DP 180 V, EP 10 V, CE 60 V) for 24:1-CoA, 1146.7/639.7 (DP 180 V, EP 10 V, CE 60 V) for 26.0-CoA, and 1144.7/637.7 (DP 180 V, EP 10 V, CE 60 V) for 26:1-CoA, respectively.

## Results

### *PpVFAD1* and *PpVFAD2* are non-essential for normal growth, but redundantly contribute to cold stress resilience

*PpVFAD1* was previously characterized (Resemann *et al*., 2021), and its desaturase activity on VLCFAs was observed by lipidomic analysis of wild type and the *vfad1* knock-out mutant grown in liquid culture. However, this phenotype presented as a deficiency, not a complete elimination, of monounsaturated VLCFAs. Further, distinct and differentiated tissues were not profiled. We therefore compared the expression profiles of *PpVFAD1* and *PpVFAD2 in silico* via the *Physcomitrella patens* eFP browser (Winter *et al*., 2007; Ortiz-Ramírez *et al*., 2016), which indicated that although *PpVFAD2* expression is consistently lower than that of *PpVFAD1*, it nevertheless is expressed, and is up-regulated in mature brown sporophytes, and in skotonema, a specialized form of protonema that grows anti-gravitropically in the dark. We examined *PpVFAD2* expression in wild type grown under our own conditions with qPCR, and confirmed low expression levels in gametophores and in mixed protonema, and six-fold elevated expression levels in dark-grown skotonema (**Figure 1A**). This suggests that *PpVFAD2* could have a specialized function, perhaps specifically supporting non-photosynthetic or anti-gravitropic growth.

**Figure 1:**
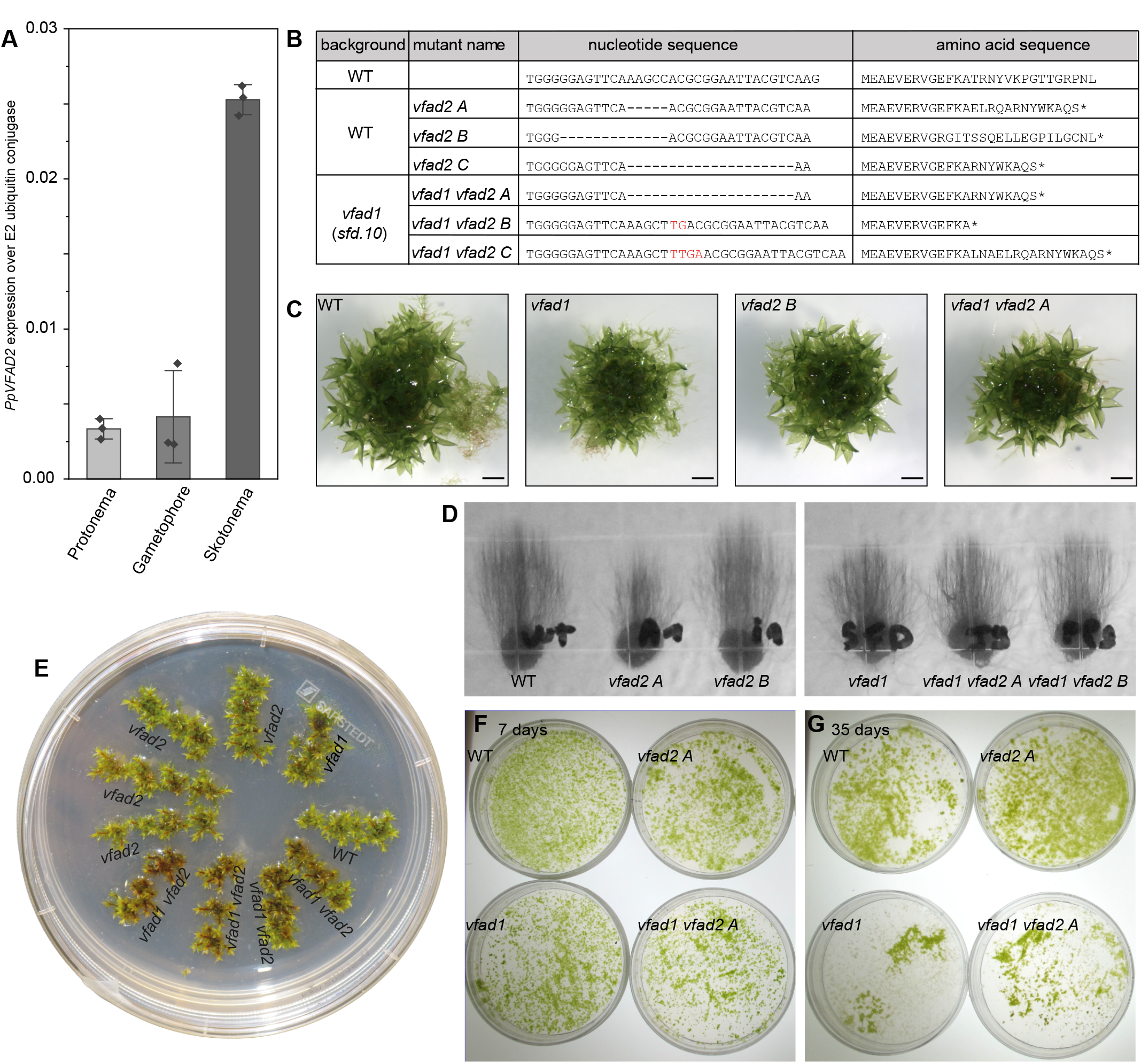
Phenotypic analysis of *Physcomitirum patens* wild type (WT), *vfad1, vfad2*, and *vfad1 vfad2*. A *PpVfAD2* expression measured by qPCR, normalized against the E2 conjugase Pp6c14_11780V6.1. Values are averages of three biological replicates, bars are standard deviation. B *vfad2* and *vfad1 vfad2* mutant lesions, which were all generated with the same CRISPR guide targeting the 5’ end of the coding sequence. The *vfad1* mutant lesion, which was generated by homologous recombination, was previously described by Resemann et al., 2021. C Four-week-old gametophores representative of each genotype. Scale bars represent 1 mm. D Skotonema cultures of representative mutants. Despite the increased expression of *VFAD2* in skotonema, the *vfad2* single mutants and *vfad1 vfad2* double mutants do not show any phenotype. E Cold treatment of gametophores. Gametophores were cultivated first for one week in control conditions, then at 6°C with for 3 weeks. F Control protonema for cold stress experiment, grown for one week in control conditions. G Cold treatment of protonema, grown for 35 days at 6°C.

We generated *vfad2* mutants using CRISPR/Cas9 as previously described (Lopez-Obando *et al*., 2016; Collonnier *et al*., 2017; Haslam *et al*., 2024), in both the wild-type background for single mutants and in *vfad1* background for *vfad1 vfad2* double mutants. Stable singles and doubles with frame-shift indels that produced premature stop codons (**Figure 1B**) were easily isolated and propagated. Both gametophore (**Figure 1C**) and skotonema tissue (**Figure 1D**) of all genotypes was cultivated for lipidomic analysis. None of the mutants showed any obvious phenotype, indicating that under our controlled growth conditions, even in skotonema where *PpVFAD2* expression is up-regulated, *PpVFAD1* and *PpVFAD2* are both non-essential.

The *A. thaliana ads2* mutant lacking monounsaturated VLCFAs was cold-sensitive (Smith *et al*., 2013), as was *P. patens vfad1* mutant protonema (Resemann *et al*., 2021). We therefore grew wild type, *vfad1, vfad2*, and *vfad1 vfad2* gametophores and protonema under cold stress conditions. Gametophores were grown first under standard conditions for one week, then transferred to cold (6°C) for three weeks. While *vfad1* and *vfad2* single mutants appeared similar to the wild type, the *vfad1 vfad2* doubles were discolored brown, and appeared less leafy and vigorous (**Figure 1E**). For cold stress of protonema, the cultures were first grown under standard conditions for one week (**Figure 1F**), after which all of the genotypes were indistinguishable. However, cultivation at 6°C substantially reduced growth of both the *vfad1* singles and the *vfad1 vfad2* doubles, whereas *vfad2* single mutants were indistinguishable from the wild type (**Figure 1G**). Altogether, these results indicate that (1) *VFAD1* and *VFAD2* are non-essential under standard non-stressed growth conditions, (2) both *VFAD1* and *VFAD2* contribute to gametophore vitality under cold stress, whereas (3) only *VFAD1* contributes to protonemal vitality under cold stress.

### *VFAD2* makes a minor contribution to VLCFA monounsaturation of sphingolipids in tissues where its expression is upregulated

We compared the sphingolipid profiles of the wild type, *vfad1, vfad2*, and *vfad1 vfad2* to establish the metabolic role of *VFAD2* alone, in comparison, or in redundancy with *VFAD1*. We profiled gametophores as a representative non-stressed vegetative growth stage (**Figure 2A-G**), as well as skotonema (**Figure 3A-G**), due to the elevated expression of *VFAD2* in this tissue type. While all sphingolipids previously detected in *P. patens* in our lab were measured, we focused on and discuss the lipids containing 24 carbon and 26 carbon VLC-acyl groups, as these consistently showed differences among genotypes. Notably, other VLCFAs present in *P. patens* sphingolipids, namely 20 and 22 carbons, could only be detected in saturated form in the wild type, suggesting that they are not preferred substrates of *VFAD1, VFAD2*, or any other desaturase in *P. patens*.

**Figure 2:**
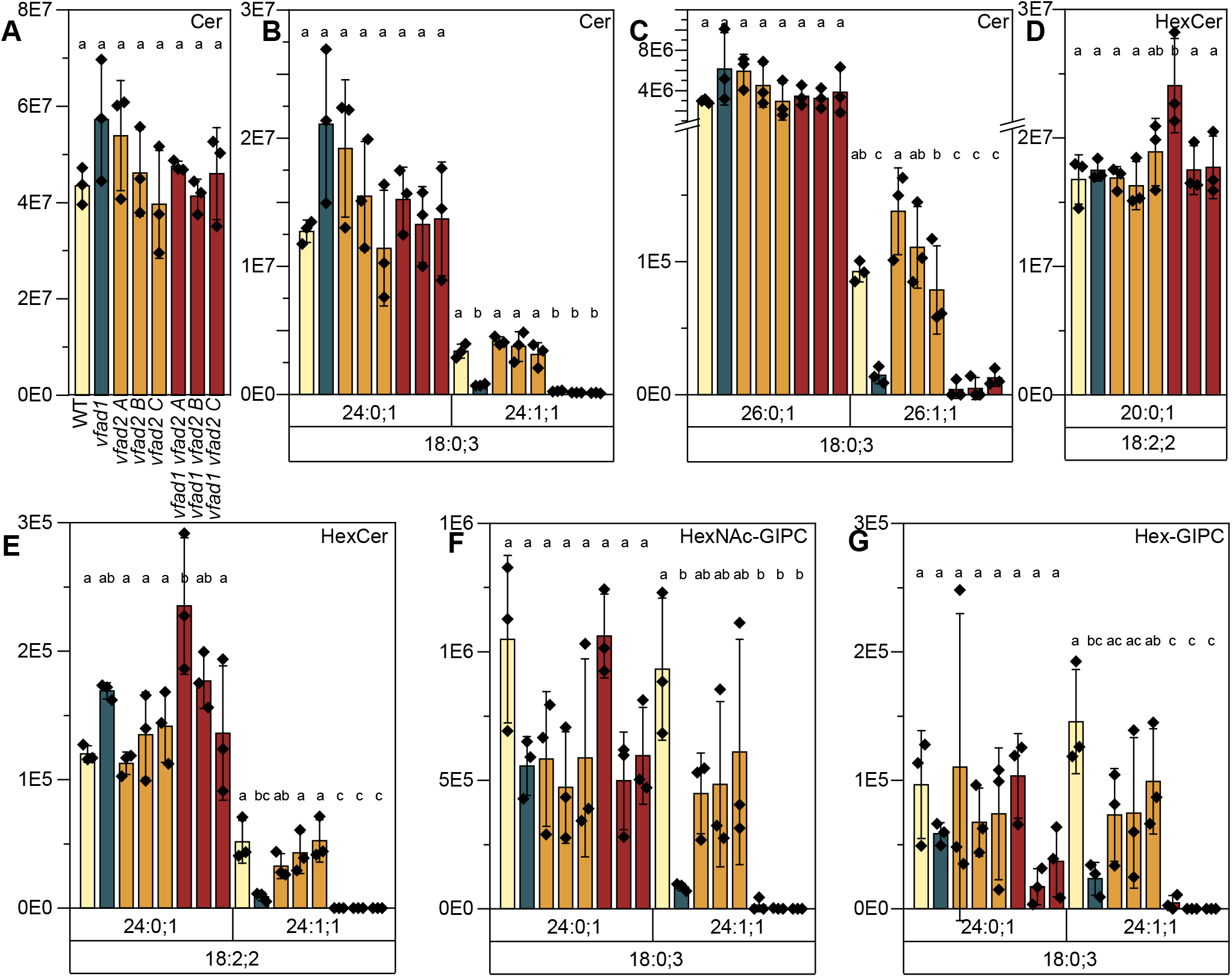
Measurement of representative, major sphingolipid species from gametophores of *vfad1, vfad2* and *vfad1 vfad2* mutants of *P. patens*. A total free ceramides, B 18:0;3/24 free ceramides, C 18:0;3/26 free ceramides, D 18:2;2/20:0;1 hexosylceramides, E 18:2;2/24 hexosylceramides, F 18:0;3/24 HexNAc-GIPCs and G 18:0;3/24 Hex-GIPCs analyzed by UPLC-nanoESI-MS/MS. All SRM peak areas were normalized to the total fatty acid methyl ester (FAMEs) amount determined for each sample. Data represent the mean ± SD of three replicates grown on separate plates. Statistical analysis was done using a one-way ANOVA with Tukey’s *post-hoc* test. Letters indicate significance at p < 0.05. Cer: free ceramide; HexCer: hexosylceramide; GIPC: glycosyl inositol phosphorylceramide.

**Figure 3:**
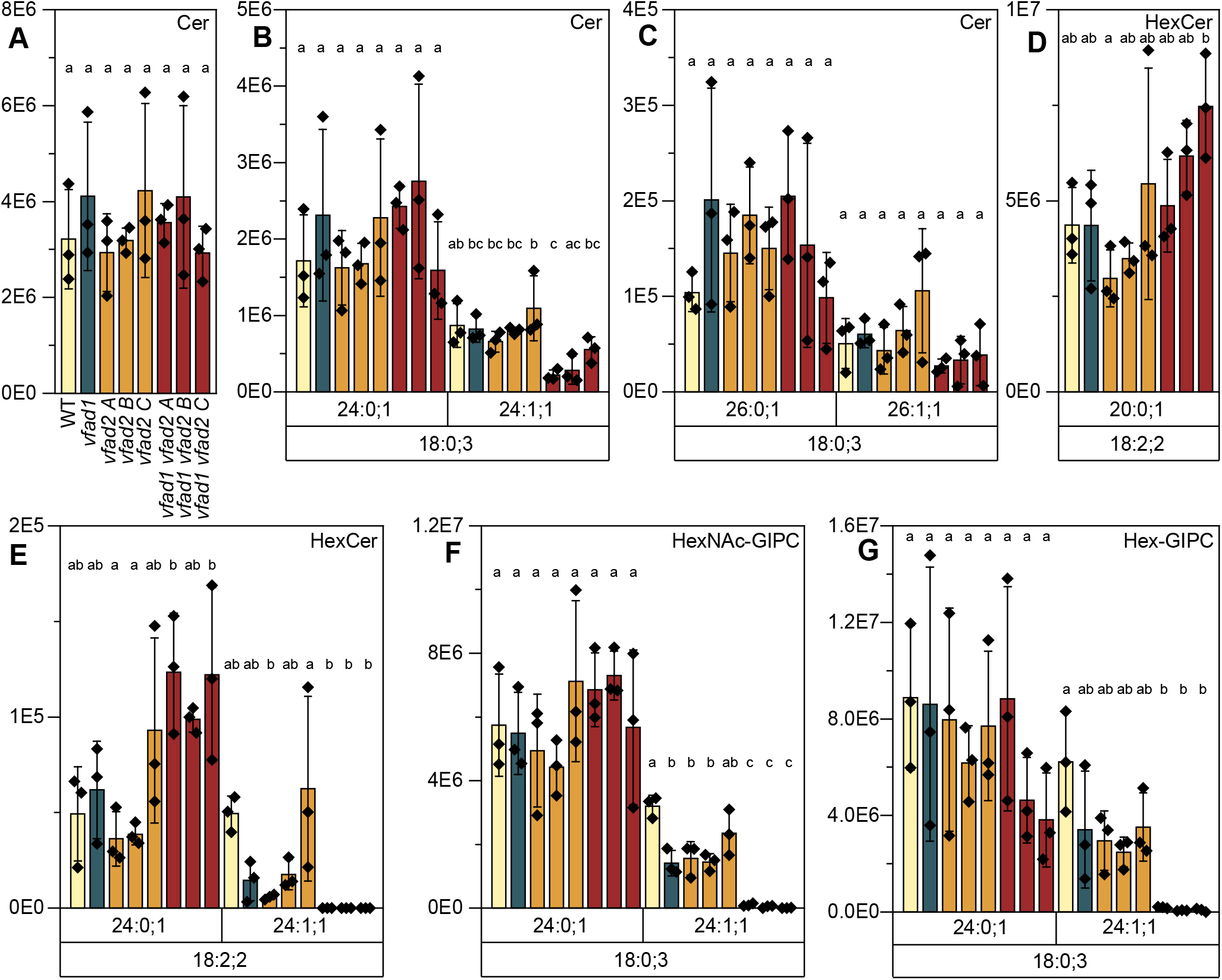
Measurement of representative, major sphingolipid species from skotonema of *vfad1, vfad2* and *vfad1 vfad2* mutants of *P. patens*. A total free ceramides, B 18:0;3/24 free ceramides, C 18:0;3/26C free ceramides, D 18:2;2/20:0;1 hexosylceramides, E18:2;2/24 hexosylceramides, F 18:0;3/24C HexNAc-GIPCs and G 18:0;3/24C Hex-GIPCs analyzed by UPLC-nanoESI-MS/MS. All SRM peak areas were normalized to the total fatty acid methyl ester (FAMEs) amount determined for each sample. Data represent the mean ± SD of three replicates grown on separate plates. Statistical analysis was done using a one-way ANOVA with Tukey’s *post-hoc* test. Letters indicate significance at p < 0.05. Cer: free ceramide; HexCer: hexosylceramide; GIPC: glycosyl inositol phosphorylceramide.

In gametophores, the total amounts of major sphingolipid classes were consistent among genotypes (**Figure 2A**), with strong shifts in molecules containing specific VLCFAs. Consistent with Resemann *et al*., 2021 we observed substantial, significant, but incomplete reductions in 24:1 components of free ceramides (**Figure 2B**), and additionally in 26:1 (**Figure 2C**) in *vfad1* single mutants. In contrast, the *vfad2* single mutants showed no consistent, significant depletion in these monounsaturated VLCFAs. The *vfad1 vfad2* double mutants also showed strong 24:1 and 26:1 free ceramide depletion, but this was not significantly different from the *vfad1* single. The total amount of hexosylceramides (HexCers) was not impacted in the mutants; the 18:2;2/20:0;1 species that makes up more than 95 % of the HexCer pool in *P. patens* was neither significantly nor consistently impacted among genotypes (Figure 2D). Low-abundant HexCers containing 24:1;1 were depleted to the same extent in *vfad1* and *vfad1 vfad2* double mutants, and not depleted in *vfad2* single mutants (Figure 2E). The same pattern of 24:1 depletion across genotypes was observed in *N*-acetyl hexosamine glycosyl inositol phosphorylceramides (HexNAc-GIPCs) (Figure 2F) and hexosyl glycosyl inositol phosphorylceramides (Hex-GIPCs) (Figure 2G). Though the depletion of monounsaturated VLCFAs in complex sphingolipids often appeared more severe in the *vfad1 vfad2* double mutants than in the *vfad1* single, this difference was insignificant.

In skotonema, we observed broadly similar trends; that is, the severe but incomplete depletion of monounsaturated VLCFA components of sphingolipids across classes, to a comparable extent in both *vfad1* and *vfad1 vfad2* (Figure 3A-G). Two notable differences in skotonema compared to gametophores were that (1) depletions in 24:1 and 26:1 fatty acids on free ceramides were insignificant and far less substantial (Figure 3B, C), and (2) depletions of 24:1 in HexNAc-GIPCs and Hex-GIPCs were more severe in *vfad1 vfad2* double mutants than in the *vfad1* single (Figure 3F, G). This more seveere reduction of monounsaturated VLCFAs in GIPCs in the double mutants is a first clear indication that *VFAD2* has a similar and redundant metabolic role to *VFAD1*, and contributes to the monounsaturation of VLCFA moieties of sphingolipids. Notably, the fact that in *vfad1* single mutants, the reduction in 24:1 was generally more severe in gametophore tissues (Figure 2E, F, and G) than in skotonema (Figure 3E, F, G), may reflect a greater contribution of *VFAD2* in skotonema which could lessen the impact of the *vfad1* single knock-out. Two surprising results of this analysis are that the *vfad1 vfad2* mutants do still produce some sphingolipids with monounsaturated VLCFAs, suggesting the existence of another desaturase contributing to this metabolic role, and (2) the retention of these residual monounsaturated VLCFAs in free ceramides, rather than incorporation in GIPCs.

### *VFAD1 and VFAD2* both contribute substantially to monounsaturation of VLCFA moieties of phosphatidylcholine and phosphatidylethanolamine

In addition to sphingolipids, the *vfad1* mutant was reported to lack phosphoglycerolipids with 24:1 fatty acyl groups, including phosphatidylcholine (PC) and phosphatidylethanolamine (PE) (Resemann *et al*., 2021). This suggests that the substrate of *VFAD1* may not necessarily be a sphingolipid as initially described. We measured PE and PC in wild type, *vfad1, vfad2*, and *vfad1 vfad2* gametophores (**Figures 4A and B**) and skotonema (**Figure 4C and D**) to determine whether *VFAD2* also impacts phosphoglycerolipid metabolism. In wild-type gametophores, we could detect PE with 24:1 and either 20:4 or 20:3 as the second fatty acyl group (**Figure 4A**), and PC with 24:1 and 20:4 fatty acids (**Figure 4B**). In both *vfad1* and *vfad2* single mutants, the average values for these genotypes showed substantial reductions compared to the wild type, up to 60 %. However, these reductions were often insignificant, especially for *vfad2*. For the *vfad1 vfad2* double mutants, the depletion was significant and more dramatic, with only trace amounts of these PE and PC species detectable in all three alleles.

**Figure 4:**
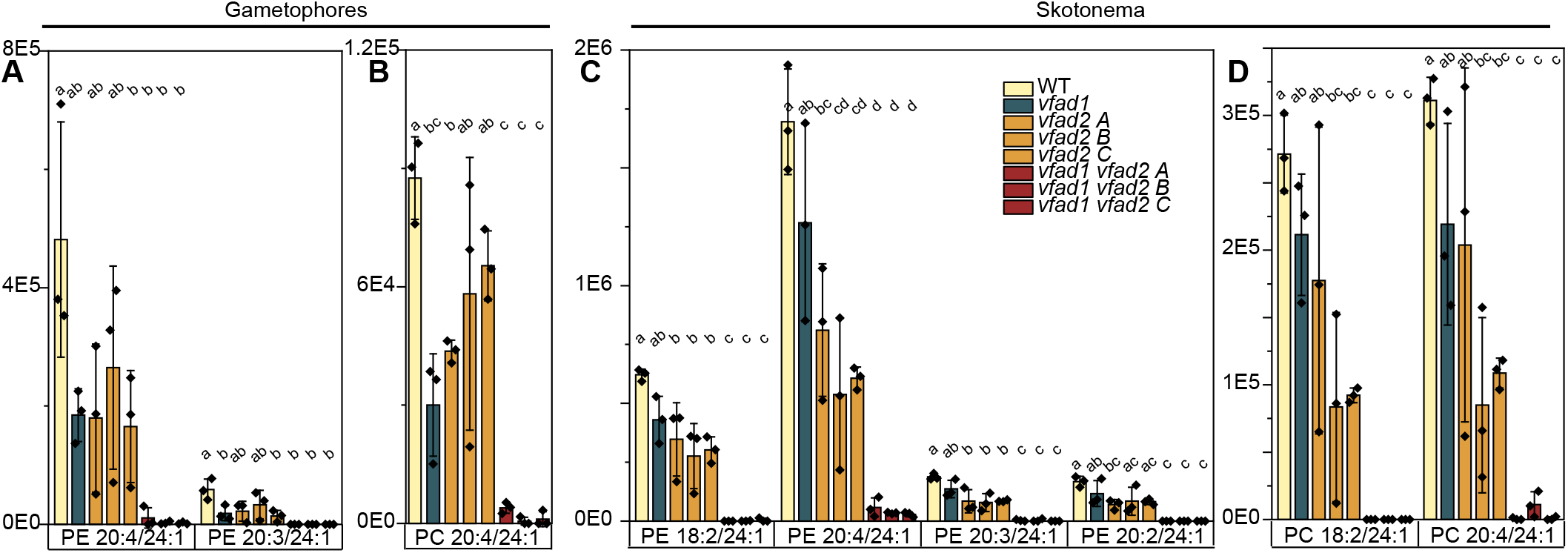
Measurement of detected phosphoglycerolipids containing 24 carbon very-long-chain fatty acids in gametophores and skotonema of *vfad1, vfad2*, and *vfad1 vfad2* mutants of *P. patens*. Phosphatidylethanolamine A, C and phosphatidylcholine B, D were measured in gametophores A, B and skotonema C, D. All unambiguous and robustly-detected lipid species containing 24 carbon VLCFAs with monounsaturation were analyzed using UPLC-nanoESI-MS/MS. The peak area of the MRMs for loss of both of the two fatty acids in each lipid species is measured and summed before being normalized to the FAMEs content of the sample. Data represent the mean ± SD of three replicates grown on separate plates. Statistical analysis was done using a one-way ANOVA with Tukey’s *post-hoc* test. Letters indicate significance at p < 0.05. PE: Phosphatidylethanolamine; PC: Phosphatidylcholine.

In wild-type skotonema, we could detect PE with 24:1 and either 18:2, 20:4, 20:3, or 20:2 fatty acids (**Figure 4C**), and PC with 24:1 and either 18:2 or 20:4 fatty acids (**Figure 4D**). The reduction in 24:1-containing PE and PC in both single mutants was modest, but more substantial for *vfad2* in this tissue type. Perhaps this reflects the increased expression level of *VFAD2* in skotonema. In contrast, 24:1-containing PE and PC were either undetectable or reduced to trace amounts in *vfad1 vfad2* double mutants.

These strong, parallel depletions in both glycerolipids and sphingolipids suggest that the substrate of VFADs could be an acyl-CoA, which would feed into both glycerolipid and sphingolipid pathways. If so, we would expect that the acyl-CoA pool would also be strongly impacted in *vfad* mutants. To examine this possibility, we attempted to extract and measure acyl-CoAs from protonema of wild-type and all three mutant genotypes. However, we could not robustly and consistently detect 24:1 in the wild-type, making meaningful comparisons to the mutants impossible.

### VFADs from both *P. patens* and *M. polymorpha* can desaturate 24 and 26 carbon VLCFAs in *S. cerevisiae*, and result in the accumulation of monounsaturated VLCFAs in sphingolipids, acyl-CoAs, and triacylglycerols

We cloned several *VFAD*s from different organisms and expressed them in the *oleic acid requiring* (*ole1*) mutant of the yeast *Saccharomyces cerevisiae. ole1* is deficient in the single gene encoding a soluble oleic acid (18:0) desaturase. The mutation is lethal, but easily rescued by supplementation with unsaturated fatty acids (Stukey *et al*., 1989, 1990). We fed yeast cells with 18:2 Δ^9,12^ to rescue the mutation, so that any monounsaturated fatty acids we detected could unambiguously be inferred to have arisen from VFAD activity, rather than elongation of 18:1. In addition to *PpVFAD1* and *PpVFAD2*, we selected gene candidates from *Marchantia polymorpha* (*MpVFAD-LIKE*), *Chlamydomonas reinhardtii* (*CrVFAD-LIKE*), and two candidates from *Thalassiosira pseudonana* (*TpVFAD-LIKE1* and *TpVFAD-LIKE2*) for expression, which we had previously identified as paralogs (Resemann *et al*., 2021).

We first measured complex sphingolipids containing 24:1 and 26:1 moieties, as sphingolipids represent the largest sink for VLCFAs (predominantly C26) in *S. cerevisiae* (Ejsing *et al*., 2009). For both the 24 and 26 chain lengths with monounsaturation, we pooled the signals for molecules with monohydroxylation, dihydroxylation, and no hydroxylations, as all of these accumulate to substantial levels in yeast cells. We could not detect any sphingolipids with monounsaturated VLCFAs in cells transformed with the empty vector, or cells expressing *CrVFAD-LIKE, TpVFAD-LIKE1*, or *TpVFAD-LIKE2*. However, cells expressing *PpVFAD1, PpVFAD2*, and *MpVFAD-LIKE* accumulated variable amounts of IPCs, MIPCs, and M(IP)_2_Cs with both 24:1 and 26:1 (**Figure 5A-C**). Accumulation of these lipids was weak in *PpVFAD2*-expressing cells (black arrows throughout **Figure 5**) compared to the other two genes, but were nevertheless consistently detected in two separate replicates of this experiment.

**Figure 5:**
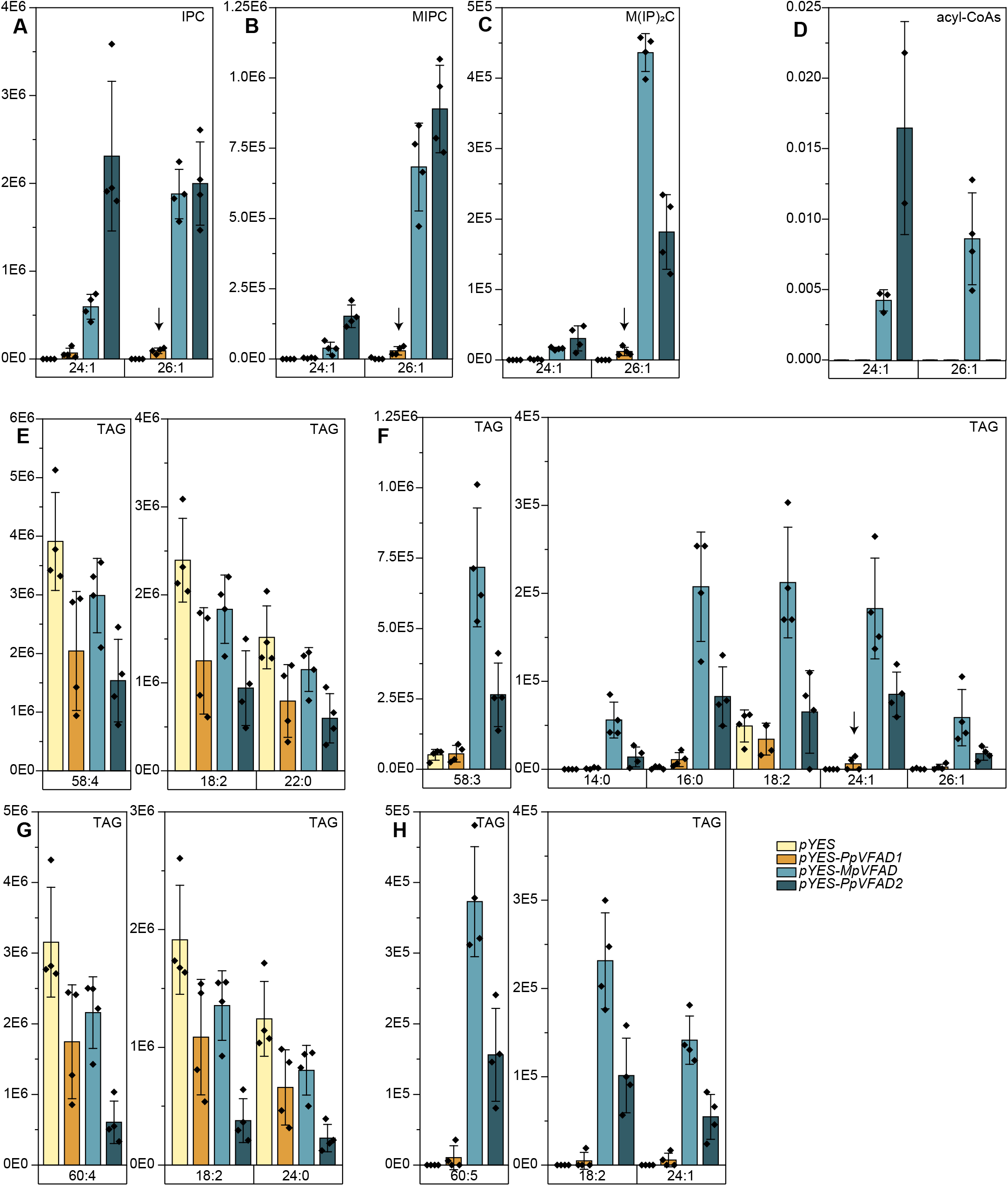
Expression of VFADs in *Saccharomyces cerevisiae* results in accumulation of monounsaturated VLCFAs in the sphingolipids A inositol phosphorylceramides (IPCs), B mannosyl inositol phosphorylceramides, and C mannosyl diinositol phosphorylceramides. These were measured by UPLC-nano-ESI-MS/MS, as the peak areas from the mass transition with neutral loss of the (IP/MIP/M(IP)_2_) headgroup. D, Measurement of monounsaturated VLC-acyl-CoAs by HPLC-ESI-MS/MS. E-H triacylglycerol (TAG) measurement by UPLC-nanoESI-MS/MS; representative species without (E, G) and with (F, H) monounsaturated VLCFAs are presented. For each, the summed peak areas of detected mass transitions (loss of each fatty acyl group) is shown on the left, and the individual transitions on the right. Bars represent the mean ± SD of four replicates, that is, independent transformed cell lines cultured separately.

We proceeded to extract and measure acyl-CoAs from cells expressing *PpVFAD1, PpVFAD2*, and *MpVFAD-LIKE* (**Figure 5D**). Both *PpVFAD1* and *MpVFAD-LIKE*-expressing cells accumulated 24:1-CoA, and *MpVFAD-LIKE* additionally accumulated 26:1-CoA. Although monounsaturated acyl-CoAs could not be detected in *PpVFAD2*, this measurement more broadly indicates that the acyl-CoA pool is impacted by VFAD activity.

We then measured glycerolipids reported to contain VLCFA moieties in *S. cerevisiae*, including phosphatidylinositol (PI) (Schneiter *et al*., 2004) and triacylglycerol (TAG) (Ejsing *et al*., 2009). We could not detect monounsaturated VLCFAs in PI, however, we did detect many TAG species that included a monounsaturated VLCFA, in the cell lines expressing *PpVFAD1* and *MpVFAD-LIKE*. Due to the volume of data and diverse TAG species detected, we selected a few representative TAG molecules (**Figure 5E-H**). This included two species that do not incorporate monounsaturated VLCFAs (**Figure 5E, G**); cells expressing VFADs generally accumulated less of these TAG species compared to the control with only the empty pYES vector. We also included two TAG species that are only possible with the inclusion of a monounsaturated (VLC)-FA (**Figure 5F, H**), as they have an odd number of double bonds. These TAGs were robustly detected in cells expressing *PpVFAD1* and *MpVFAD-LIKE*, whereas only trace amounts of either were detected in the pYES control and cells expressing *PpVFAD2*. Altogether, expression of *VFAD*s in *S. cerevisiae* demonstrated that these genes impact not only sphingolipid metabolism, but also the acyl-CoA profile and the profile of neutral storage lipids.

## Discussion

### Both *ADS*s and *VFAD*s have patchy distribution patterns among embryophytes

We previously identified and characterized *P. patens* VFAD1/SFD, a front-end, cytochrome b5 fusion desaturase that has an analogous metabolic role to *A. thaliana* ADS2, a methyl-end desaturase. In spite of their distinct domain structures and phylogeny, both enzymes contribute to the monounsaturation of VLCFA moieties of sphingolipids as well as glycerolipids. They also have convergent functionality in that they contribute to cold stress adaptation: both *Atads2* and *Ppvfad1* mutants show reduced fitness under prolonged exposure to cold stress, and ectopic expression of *PpVFAD1* can complement the cold sensitivity of *Atads2* mutants. Despite our efforts to investigate *PpVFAD2* activity in skotonema where its transcript levels are elevated, we observed no change in growth or development in this tissue type in *Ppvfad2* mutants, nor in *Ppvfad1 vfad2* double mutants. Indeed, the only indication of a physiological role of *PpVFAD2* was the intensified signs of cold stress in *Ppvfad1 vfad2* double mutant gametophores compared to both wild-type and *Ppvfad1*.

Notably, neither ADS-type nor VFAD-type desaturases are ubiquitous in embryophytes. ADS-related enzymes are dispersed throughout eukaryotes, are well-characterized in yeast and mammalian systems, and have been identified in diverse plants ranging from *Picea glauca* (white spruce) (Marillia *et al*., 2002), *Limnanthes alba* (white meadowfoam) (Pollard and Stumpf, 1980; Moreau *et al*., 1981; Cahoon *et al*., 2000), and *Anemone leveillei* (woodland windflower) (Sayanova *et al*., 2007). However, there are no homologs of ADS in rice, brachypodium, or maize (Smith *et al*., 2013), and no unsaturated VLCFA moieties of sphingolipids have been detected in rice (Ishikawa *et al*., 2016). Collectively, this hints that plant ADSs, though generally widespread in eukaryotes and seemingly so among most tracheophytes, could have been lost in the monocot lineage. More thorough sampling of monocot species and functional characterization is necessary to test this hypothesis.

Phylogenetic analysis of VFADs throughout eukaryotes also revealed widespread but patchy distribution of these enzymes, with candidate genes identified among lineages as diverse as diatoms, choanoflagellates, and chytrid fungi. Yet, among a diverse sampling of the embryophytes, VFADs could only be identified in *P. patens* and *M. polymorpha* - both setaphytes. VFAD as a derivative of front-end, cytb5-fusion enzymes fits with the conservation of the LC-PUFA pathway, as these enzymes play a central role in lipid metabolism in bryophytes. Overall, the uneven distribution of ADS2-LIKEs and VFAD-LIKEs prompts speculation as to whether similar metabolic activity exists in some form in the remaining plant lineages - i.e. monocots, hornworts, lycophytes, monilophytes - and if so, what enzymes are responsible for such activity.

### Residual monounsaturated VLCFAs accumulate in different lipid classes in *vfad* mutants

Comparing the *vfad1, vfad2*, and *vfad1 vfad2* metabolic phenotypes, we observed surprising trends in which lipids were most severely impacted by mutations. In the single *vfad1* mutant gametophores, with only partial reductions in 24:1-and 26:1-containing lipids, there was a more severe impact on sphingolipids than glycerolipids. (In skotonema, the lipid phenotypes of *vfad1* single mutants were, with the sole exception of 24:1:1 HexNAc-GIPCs, non-significant). This could either reflect a metabolic prioritization of glycerolipid synthesis over sphingolipid synthesis, or alternatively, the accessibility of VFAD1 product to either pathway. Reductions in *vfad2* single mutants were weak; however, the double *vfad1 vfad2* mutants were surprising in that the most robust signals for residual monounsaturated VLCFAs were detected in the free ceramides of skotonema.

The source of residual monounsaturated VLCFAs in *vfad1 vfad2* remains unknown. The simplest explanation is that they could be synthesized by elongation of monounsaturated LCFAs produced by the soluble, plastidial acyl-ACP desaturase. Alternatively, we reason that the metabolic source of monounsaturated VLCFAs in the double *vfad1 vfad2* mutant should produce these in a way that, in contrast to VFADs, results in preferential use in sphingolipid metabolism. We suggest that the Δ8-LCB desaturase, which is also localized to the ER and expected to accept LCBs conjugated to free ceramides as substrates, could also have some promiscuous functionality with the fatty acyl moieties of free ceramides. The products could, subsequently and inefficiently, be distributed to other lipid pools, via release by ceramidases and incorporation into glycerolipids. This hypothesis could be tested by isolation and lipidomic analysis of *Ppsd8d vfad1 vfad2* triple mutants.

### Acyl-CoAs are most likely the substrates of both ADSs and VFADs

Characterization of AtADS2 demonstrated that its loss affected sphingolipid, phosphoglycerolipid, and acyl-CoA profiles (Smith *et al*., 2013). Similarly, loss of PpVFAD1 and PpVFAD2 affected sphingolipid and phosphoglycerolipid profiles. While we were unable to detect VLC monounsaturated acyl-CoAs in *P. patens* tissue, lipid profiling of *S. cerevisiae* heterologously-expressing *PpVFAD1* and *M. polymorpha MpVFAD-LIKE* demonstrated that they are sufficient to produce 24:1 and 26:1 acyl-CoAs. The yeast cells also accumulated 24:1 and 26:1 in TAG, and in sphingolipids. The simplest model to explain the impact of both ADSs and VFADs on acyl-CoAs, glycerolipids, and sphingolipids is that their substrate is an acyl-CoA, which could directly feed into glycerolipid and sphingolipid assembly. As with cold-stress resilience imparted by both ADSs and VFADs, their (inferred) metabolic similarity in modifying acyl-CoAs is striking given their distinct domain structure and phylogeny. This again highlights a disconnect between classification and nomenclature of desaturases based on phylogeny vs. their *in vivo* physiological and metabolic functions, and urges caution in the nomenclature of newly-identified enzymes.

## Supplemental Data

Supplemental Figure 1: Complete ceramide backbone profiles for sphingolipid measurements of skotonema of *vfad1, vfad2*, and *vfad1 vfad2* mutants of *P. patens*.

Supplemental Table 1: Primers used in the study.

## Acknowledgements

We thank Dr. Ellen Hornung and Sabine Freitag for technical support and contributions to this project, Dr. Patricia Scholz for the R script used for statistical analysis, and Dr. Alisa Keyl for providing the *Marchantia polymorpha* thalli cDNA.

## CRediT Author Contributions

TMH, IF: Conceptualization, Funding Acquisition, Project Administration, Supervision; PD, CH, TMH: Data Curation, Investigation, Validation; PD, TMH: Visualization, Writing - Original Draft; CH, IF: Resources; PD, CH, TMH, IF: Formal Analysis, Methodology, Writing - Review and Editing

## Abbreviations

ACP: acyl carrier protein
ADS: ACYL-COENZYME A DESATURASE
CoA: Coenzyme A
FAD: FATTY ACYL DESATURASE
GIPC: glycosyl inositol phosphorylceramide
MGDG: monogalactosyldiacylglycerol
OLE: OLEIC ACID-REQUIRING
PC: phosphatidylcholine
PE: phosphatidylethanolamine
PM: plasma membrane
SFD: SPHINGOLIPID FATTY ACID DESATURASE (renamed VFAD)
TAG: triacylglycerol
VFAD: VERY-LONG-CHAIN FATTY ACYL DESATURASE
VLCFA: very-long-chain fatty acid

## Literature Cited

Arondel V, Lemieux B, Hwang I, Gibson S, Goodman HM, Somerville CR. 1992. Map-Based Cloning of a Gene Controlling Omega-3 Fatty Acid Desaturation in Arabidopsis. Science 258, 1353–1355.

Le Bail A, Scholz S, Kost B. 2013. Evaluation of Reference Genes for RT qPCR Analyses of Structure-Specific and Hormone Regulated Gene Expression in Physcomitrella patens Gametophytes. PLOS ONE 8, e70998.

Browse J, Kunst L, Anderson S, Hugly S, Somerville C. 1989. A Mutant of Arabidopsis Deficient in the Chloroplast 16:1/18:1 Desaturase. Plant Physiology 90, 522–529.

Browse J, McConn M, James D, Miquel M. 1993. Mutants of Arabidopsis deficient in the synthesis of α-linolenate: Biochemical and genetic characterization of the endoplasmic reticulum linoleoyl desaturase. Journal of Biological Chemistry 268, 16345–16351.

Browse J, McCourt P, Somerville CR. 1985. A Mutant of Arabidopsis Lacking a Chloroplast-Specific Lipid. Science 227, 763–765.

Browse J, McCourt P, Somerville C. 1986. A Mutant of Arabidopsis Deficient in C18:3 and C16:3 Leaf Lipids. Plant Physiology 81, 859–864.

Cahoon EB, Marillia EF, Stecca KL, Hall SE, Taylor DC, Kinney AJ. 2000. Production of fatty acid components of meadowfoam oil in somatic soybean embryos. Plant Physiology 124, 243–251.

Chen M, Thelen JJ. 2013. ACYL-LIPID DESATURASE2 is required for chilling and freezing tolerance in Arabidopsis. Plant Cell 25, 1430–1444.

Collonnier C, Epert A, Mara K, et al. 2017. CRISPR-Cas9-mediated efficient directed mutagenesis and RAD51-dependent and RAD51-independent gene targeting in the moss Physcomitrella patens. Plant Biotechnology Journal 15, 122–131.

Domergue F, Abbadi A, Zähringer U, Moreau H, Heinz E. 2005. In vivo characterization of the first acyl-CoA Δ6-desaturase from a member of the plant kingdom, the microalga Ostreococcus tauri. Biochemical Journal 389, 483–490.

Ejsing CS, Sampaio JL, Surendranath V, Duchoslav E, Ekroos K, Klemm RW, Simons K, Shevchenko A. 2009. Global analysis of the yeast lipidome by quantitative shotgun mass spectrometry. Proceedings of the National Academy of Sciences of the United States of America 106, 2136–2141.

Fernandez-Pozo N, Haas FB, Meyberg R, et al. 2020. PEATmoss (Physcomitrella Expression Atlas Tool): a unified gene expression atlas for the model plant Physcomitrella patens. Plant Journal 102, 165–177.

Garcés R, Mancha M. 1993. One-step lipid extraction and fatty acid methyl esters preparation from fresh plant tissues. Analytical Biochemistry 211, 139–143.

Gibson S, Arondel V, Iba K, Somerville C. 1994. Cloning of a temperature-regulated gene encoding a chloroplast ω-3 desaturase from Arabidopsis thaliana 1. Plant Physiology 106, 1615–1621.

Haslam TM, Herrfurth C, Feussner I. 2024. Diverse INOSITOL PHOSPHORYLCERAMIDE SYNTHASE mutant alleles of Physcomitrium patens offer new insight into complex sphingolipid metabolism. New Phytologist 242, 1189–1205.

Haslam RP, Larson TR. 2021. Techniques for the Measurement of Molecular Species of Acyl-CoA in Plants and Microalgae. Plant Lipids: Methods and Protocols. 203–218.

Heilmann I, Pidkowich MS, Girke T, Shanklin J. 2004. Switching desaturase enzyme specificity by alternate subcellular targeting. Proceedings of the National Academy of Sciences of the United States of America 101, 10266–10271.

Herrfurth C, Liu Y-T, Feussner I. 2021. Targeted Analysis of the Plant Lipidome by UPLC-NanoESI-MS/MS. In: Bartels D, Dörmann P, eds. Plant Lipids: Methods and Protocols. Springer Nature, 135–155.

Ishikawa T, Ito Y, Kawai-Yamada M. 2016. Molecular characterization and targeted quantitative profiling of the sphingolipidome in rice. Plant Journal 88, 681–693.

Kazaz S, Miray R, Lepiniec L, Baud S. 2022. Plant monounsaturated fatty acids: Diversity, biosynthesis, functions and uses. Progress in Lipid Research 85.

Kunst L, Browse J, Somerville C. 1989. A mutant of arabidopsis deficient in desaturation of palmitic acid in leaf lipids. Plant Physiology 90, 943–947.

Larson TR, Edgell T, Byrne J, Dehesh K, Graham IA. 2002. Acyl CoA profiles of transgenic plants that accumulate medium-chain fatty acids indicate inefficient storage lipid synthesis in developing oilseeds. The Plant Journal 32, 519–527.

Larson TR, Graham IA. 2001. A novel technique for the sensitive quantification of acyl CoA esters from plant tissues. The Plant Journal 25, 115–125.

Liu YC, Vidali L. 2011. Efficient polyethylene glycol (PEG) mediated transformation of the moss Physcomitrella patens. Journal of Visualized Experiments, 2–5.

Lopez-Obando M, Hoffmann B, Géry C, Guyon-Debast A, Téoulé E, Rameau C, Bonhomme S, Nogué F. 2016. Simple and efficient targeting of multiple genes through CRISPR-Cas9 in Physcomitrella patens. G3: Genes, Genomes, Genetics 6, 3647–3653.

Marillia EF, Giblin EM, Covello PS, Taylor DC. 2002. A desaturase-like protein from white spruce is a Δ9 desaturase. FEBS Letters 526, 49–52.

Maronova M, Kalyna M. 2016. Generating Targeted Gene Knockout Lines in Physcomitrella patens to Study Evolution of Stress-Responsive Mechanisms. Methods Mol Biol. 1398, 221–234.

McConn M Le, Hugly S, Browse J, Somerville C. 1994. A mutation at the fad8 locus of Arabidopsis identifies a second chloroplast ω-3 desaturase. Plant Physiology 106, 1609–1614.

Miquel M, Browse J. 1992. Arabidopsis mutants deficient in polyunsaturated fatty acid synthesis: Biochemical and genetic characterization of a plant oleoyl-phosphatidylcholine desaturase. Journal of Biological Chemistry 267, 1502–1509.

Moreau RA, Pollard MR, Stumpf PK. 1981. Properties of a Δ5-Fatty Acyl-CoA Desaturase in the Cotyledons of Developing Limnanthes alba. Archives of Biochemistry and Biophysics 209, 376–384.

Napier JA, Michaelson L V., Stobart AK. 1999. Plant desaturases: Harvesting the fat of the land. Current Opinion in Plant Biology 2, 123–127.

Ohlrogge JB, Browse J. 2011. Lipid Biosynthesis. Society 7, 957–970.

Ortiz-Ramírez C, Hernandez-Coronado M, Thamm A, Catarino B, Wang M, Dolan L, Feijó JAA, Becker JDD. 2016. A Transcriptome Atlas of Physcomitrella patens Provides Insights into the Evolution and Development of Land Plants. Molecular Plant 9, 205–220.

Pollard MR, Stumpf PK. 1980. Biosynthesis of C 20 and C 22 Fatty Acids by Developing Seeds of Limnanthes alba. Plant Physiology 66, 649–655.

Resemann HC, Herrfurth C, Feussner K, et al. 2021. Convergence of sphingolipid desaturation across over 500 million years of plant evolution. Nature Plants 7, 219–232.

Saavedra L, Catarino R, Heinz T, Heilmann I, Bezanilla M, Malhó R. 2015. Phosphatase and tensin homolog is a growth repressor of both rhizoid and gametophore development in the moss Physcomitrella patens. Plant Physiology 169, 2572–2586.

Sayanova O, Haslam R, Caleron MV, Napier JA. 2007. Cloning and characterization of unusual fatty acid desaturases from Anemone leveillei: Identification of an acyl-coenzyme A C20 Δ5-desaturase responsible for the synthesis of sciadonic acid. Plant Physiology 144, 455–467.

Schneiter R, Brügger B, Amann CM, Prestwich GD, Epand RF, Zellnig G, Wieland FT, Epand RM. 2004. Identification and biophysical characterization of a very-long-chain-fatty-acid-substituted phosphatidylinositol in yeast subcellular membranes. Biochemical Journal 381, 941–949.

Schwarz P, Herrfurth C, Steinem C, Feussner I. 2022. Lipidomics of Thalassiosira pseudonana as a function of valve SDV synthesis. Journal of Applied Phycology 34, 1471–1481.

Shanklin J, Cahoon EB. 1998. Desaturation and related modifications of fatty acids. Annual Review of Plant Biology 49, 611–641.

Shanklin J, Somerville C. 1991. Stearoyl-acyl-carrier-protein desaturase from higher plants is structurally unrelated to the animal and fungal homologs. Proceedings of the National Academy of Sciences of the United States of America 88, 2510–2514.

Smith MA, Dauk M, Ramadan H, Yang H, Seamons LE, Haslam RP, Beaudoin F, Ramirez-Erosa I, Forseille L. 2013. Involvement of Arabidopsis acyl-coenzyme A desaturase-like2 (At2g31360) in the biosynthesis of the very-long-chain monounsaturated fatty acid components of membrane lipids. Plant Physiology 161, 81–96.

Sperling P, Heinz E. 2001. Desaturases fused to their electron donor. European Journal of Lipid Science and Technology 103, 158–180.

Sperling P, Ternes P, Zank TK, Heinz E. 2003. The evolution of desaturases. Prostaglandins, Leukotrienes and Essential Fatty Acids 68, 73–95.

Stukey JE, McDonough VM, Martin CE. 1989. Isolation and characterization of OLE1, a gene affecting fatty acid desaturation from Saccharomyces cerevisiae. Journal of Biological Chemistry 264, 16537–16544.

Stukey JE, McDonough VM, Martin CE. 1990. The OLE1 gene of Saccharomyces cerevisiae encodes the delta 9 fatty acid desaturase and can be functionally replaced by the rat stearoyl-CoA desaturase gene. Journal of Biological Chemistry 265, 20144–20149.

Tarazona P, Feussner K, Feussner I. 2015. An enhanced plant lipidomics method based on multiplexed liquid chromatography – mass spectrometry reveals additional insights into cold- and drought-induced membrane remodeling. The Plant Journal 84, 621–633.

Winter D, Vinegar B, Nahal H, Ammar R, Wilson G V., Provart NJ. 2007. An ‘electronic fluorescent pictograph’ browser for exploring and analyzing large-scale biological data sets. PLoS ONE 2, e718.

